# Transient fetal type 2 bovine viral diarrhea virus infection protects beef cattle from postnatal challenge with type 1b bovine viral diarrhea virus and *Mannheimia haemolytica*

**DOI:** 10.64898/2026.11.23.768602

**Authors:** Hana Van Campen, Huey Yi Loh, Octavio Guimaraes, Jessica N. Kincade, Morgan Adkins, Jordan M. Eder, Thomas R. Hansen, Terry E. Engle

## Abstract

Bovine respiratory disease (BRD) is the most significant disease in feedlot cattle, impacting animal growth, performance, and profitability. Acute bovine viral diarrhea virus (BVDV) infections have been shown to increase the occurrence of BRD. Fetal infection with BVDV during late gestation results in transient fetal infections (TI). At birth, TI calves are negative for BVDV but positive for BVDV-specific antibodies. The objective of this experiment was to determine the effect of TI status on the susceptibility to BRD and the subsequent rate of gain in feedlot cattle. Eight-month-old type 2 BVDV TI and uninfected control heifers (TI-BRD n=5; CON-BRD n=6) were challenged with an intranasal inoculation of non-cytopathogenic (ncp) type 1b BVDV, followed by an intratracheal inoculation with *Mannheimia haemolytica*. TI and uninfected heifers (TI-CON n=6; CON-CON n=5) were sham-inoculated with media and phosphate buffered saline (PBS) to serve as BRD challenge controls. Heifers were weighed at the beginning and end of the experiment, were evaluated daily for clinical signs of respiratory disease, and jugular blood samples were collected on d 0, 3, 4, 6, 8, 10, 17, and 31 post-BVDV inoculation. Complete blood count (CBC), flow cytometry, BVDV RTPCR, and serum neutralizing antibody titers for type 1b and type 2 BVDV were measured in blood and serum samples. Average daily gain was similar across treatments. Fetal infection with ncp type 2 BVDV in late gestation resulted in cross-protective immunity, which reduced (P < 0.05) clinical scores, lymphopenia, and viremia in TI heifers following postnatal challenge with a noncytopathic type 1b BVDV and *Mannheimia haemolytica* compared to challenged control heifers. These findings indicate that fetal BVDV TI may protect offspring from the deleterious effects of combined type 1b BVDV and *Mannheimia haemolytica* infection during the postnatal period.

**Lay summary:** A pregnant cow infected with bovine viral diarrhea virus (BVDV) during late gestation results in a transient fetal BVDV infection (TI). At birth, TI calves are negative for BVDV but positive for BVDV-specific antibodies. BVDV infections have been shown to increase the susceptibility of cattle to bovine respiratory disease (BRD). To investigate the effect of fetal TI on susceptibility or resistance to BRD, 8-month-old type 2 BVDV TI and control (CON) heifers were challenged with BVDV (type 1b), followed by inoculation of a BRDC-associated bacterium or sham-control. Heifers were evaluated daily for clinical signs, and blood samples were collected for a complete blood count and flow cytometry analysis. The presence of BVDV in blood and antibodies to type 1b and type 2 BVDV were measured. Change in body weight and average daily gain (ADG) over the 31-day experiment was determined for all heifers. Antibodies to type 2 BVDV in the TI heifers provided cross-protective immunity that reduced the severity of clinical signs, the amount of BVDV in blood, leukopenia, and lymphopenia following postnatal challenge with type 1b BVDV and *Mannheimia haemolytica* compared to control heifers. Body weight and overall ADG were similar between treatment groups during the experimental period.

**Teaser Text:** Heifer calves born to heifers infected with type 2 BVDV on d 175 of gestation are protected from severe disease, fever, viremia, leukopenia and lymphopenia when challenged with type1b BVDV and *M. haemolytica* at 8 months of age.

**Graphical abstract:** 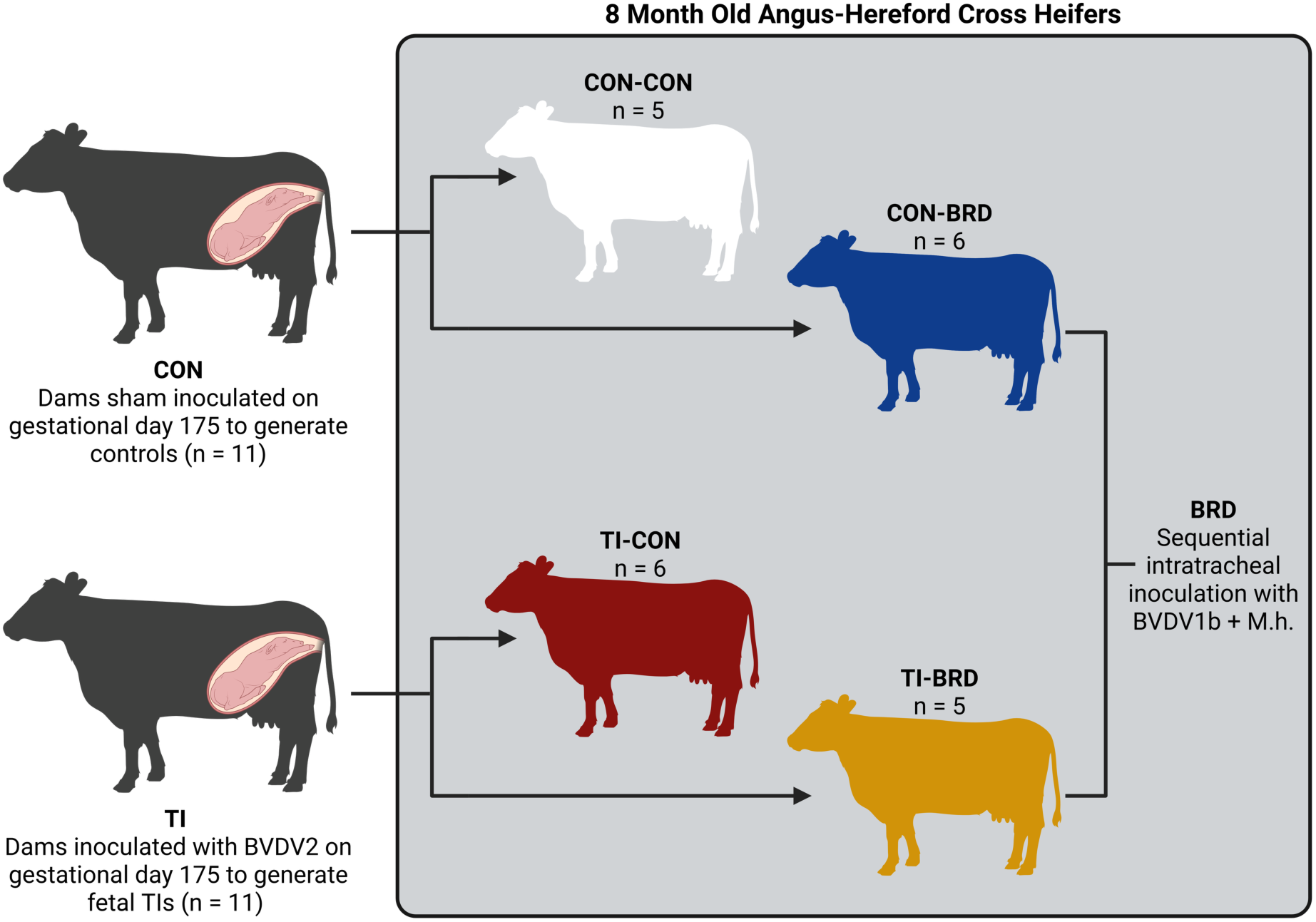

## INTRODUCTION

Bovine respiratory disease (BRD) is the most significant disease complex in feedlot cattle, impacting feedlot cattle performance and profitability (Loneragan et al., 2001; Holland et al., 2010; Cernicchiaro et al., 2013). It has been estimated to cost the American feedlot industry between $600 and $800 million annually due to decreased animal production and increased veterinary treatment costs (Chirase and Greene, 2001; USDA, 2011). Many factors contribute to BRD, including weaning and shipping stress, weather, age, weight of calves on arrival, and nutritional and vaccination status (Taylor et al., 2010). Many viruses and bacteria have been identified as being associated with BRD, including bovine viral diarrhea virus (BVDV) and *Mannheimia haemolytica* (M. haemolytica; Ruiz et al., 2022; Fritzen et al., 2024; David and Abuelo, 2025). *M. haemolytica* is frequently isolated from cattle with BRD and is considered a major contributor to lung lesions (Cozens et al., 2019). Animals that were infected with BVDV experienced respiratory and gastrointestinal disease and had a higher risk of BRD development, as BVDV also plays a role in suppressing the immune system, which allows superinfections by other respiratory pathogens (Brodersen, 2014; Gaudino et al., 2022).

Vertical transmission with noncytopathic (ncp) BVDV before day 125 of gestation, prior to full development of the fetal immune system, results in the birth of immunotolerant persistently infected (PI) calves (Brownlie et al., 1984; McClurkin et al., 1984); whereas, fetal infection after more advanced development of the fetal immune system (infection of the dam at ∼150 days), results in clearance of BVDV from the fetus and the birth of seropositive calves but BVDV negative, referred to as congenitally infected or transiently infected (TI) calves (Muñoz-Zanzi et al., 2003). Transiently infected calves are prone to a variety of health problems (Muñoz-Zanzi et al., 2003) and were suspected to be lighter in weight than uninfected calves in their cohort (Waldner and Kennedy, 2008). The effects of TI cattle on their health in the feedlot have only recently been investigated (Engle et al., 2024). Experimentally produced TI calves were lighter in weight, had an impaired ability to digest feed, and a reduced rate of gain compared to control calves, leading to increased costs and reduced profitability (Engle et al., 2024). The negative effects of fetal TI status on nutrient digestibility, rumen fermentation characteristics, and blood biomarkers of inflammation suggest a dysregulation of the immune system (Engle et al., 2024). This relationship between the immune system and inflammation is supported by peripheral white blood cell (WBC) methylome data indicating that epigenetic changes occur in multiple genes impacting immune system pathways in TI cattle (Van Campen et al., 2024; Kincade et al., 2025). Pathway analyses of hyper-and hypomethylated genes in TI WBC DNA suggest both inhibitory and stimulatory effects on the immune systems of TI cattle compared to uninfected controls. However, limited to no research has investigated the relationship between the epigenetically modified genes of the immune system of TI calves and their susceptibility or resistance to BRD. Therefore, we proposed to test the null hypothesis that TI cattle will have the same susceptibility or resistance to BRD as uninfected cattle. The objective of this experiment was to determine the effect of TI status on susceptibility or resistance to BRD and the subsequent rate of gain in feedlot cattle.

## MATERIALS AND METHODS

### Ethical statement

Prior to the initiation of this experiment, all animal care, handling, and procedures described herein were approved by the Colorado State University (CSU) Animal Care and Use Committee (IACUC Protocol approval number: 1656).

### Animals

An equal number of TI (n = 11) and uninfected control (CON; n = 11) Hereford x Angus heifers sired by the same bull were generated by inoculating pregnant heifers with a noncytopathic (ncp) type 2 BVDV (TI) or sham control media (CON) on gestational day 175 as previously described in Engle et al. (2024) and Van Campen et al. (2024). At 7 months of age, all heifers (n = 22) were weaned and transported to feedlot pens at the Agricultural Research and Education Center (ARDEC) at CSU. For a detailed description of animal management of cattle in the research feedlot, see Engle et al. (2024). Briefly, upon arrival, cattle were weighed, implanted with a growth promotant (Synovex-H; Zoetis, Kalamazoo, MI), vaccinated (Inforce 3, Zoetis, Kalamazoo, MI), dewormed (Cydectin®, Zoetis, Kalamazoo, MI), and adjusted to a standard high-concentrate steam-flaked corn-based finishing diet over a 28-day period.

### Bovine respiratory disease inocula preparation

The ncp type 1b BVDV used in the BRD challenge was isolated from a blood sample from a PI calf in a group of weaned commercial calves that experienced mortality due to pneumonia. The ncp type 1b BVDV stock was propagated in bovine turbinate (BT) cells in Dulbecco’s minimal essential medium (DMEM) + 2% horse serum (HS), and the tissue culture infectious dose (TCID)_50_/ml was determined as previously described (Smirnova et al., 2008). The ncp type 1b BVDV inoculum contained 4.0 log_10_ TCID_50_/ml.

An isolate of *M. haemolytica* grown from a feedlot calf with pneumonia was obtained from the Bacteriology Section of CSU Veterinary Diagnostic Laboratory (CSU-VDL; accession #F2000-99372). The *M. haemolytica* isolate was positive for leukotoxin 1a as determined by PCR (Washington Animal Disease Diagnostic Laboratory, accession #16189). The isolate was sensitive to ampicillin, ceftiofur, chlortetracycline, danofloxacin, enrofloxacin, florfenicol, gentamicin, oxytetracycline, spectinomycin, tilmicosin, trimethoprim-sulfamethoxazole, and tulathromycin. The inoculum contained 6 x 10^9^ colony-forming units (CFU) of *M. haemolytica* in 10 ml of PBS (Absorbance of 1.083 to 1.404 at 290 nm; Equivalent to 1.5 to 2.0 x 10^10^ CFU/ml (Mosier et al., 1998; Burciaga-Robles et al., 2010).

### BRD challenge experimental design

At 8 months of age, 11 TI and 11 CON heifers were randomly assigned, within TI or CON groups, to the control (CON) or BRD challenge groups (BRD), respectively. A total of 4 treatment groups were generated in this study: 1) CON-CON: uninfected heifers received 2 ml of media (DMEM + 2% HS) instilled into each nostril (total of 4 ml) on d 0, followed by intratracheal instillation of 10 ml of phosphate-buffered saline (PBS) on d 3 (n = 5); 2) TI – CON: TI heifers received 2 ml of media (DMEM + 2% HS) instilled into each nostril (total of 4 ml) on d 0, followed by intratracheal instillation of 10 ml of PBS on d 3 (n=6); 3) CON – BRD: uninfected heifers received 2 ml of 4.0 log_10_ TCID_50_ of the ncp type 1b BVDV in DMEM + 2% HS instilled into each nostril (total of 4 ml) on d 0, followed by intratracheal instillation of *M. haemolytica* on d 3 (n = 6); and 4) TI – BRD: TI heifers received 2 ml of 4.0 log_10_ TCID_50_ of the ncp type 1b BVDV in DMEM + 2% HS instilled into each nostril (total of 4 ml) on d 0, followed by intratracheal instillation of *M. haemolytica* on d 3 (n = 5). The BRD-challenged heifers were housed separately from the controls to prevent the transmission of BRD inoculum components (ncp type 1b BVDV and *M. haemolytica*) between groups. Any heifer with a rectal temperature ≥ 40.6 °C was treated with Florfenicol per label instructions.

### Clinical scores, rectal temperatures, respiration rates, and body weights

All heifers were weighed on day (d 0) of the BRD challenge and 31 d later. Following inoculation (d 0), heifers were observed for behavioral changes (score of 0 to 4: 0 = no discomfort and normal behavior, 1 = mild discomfort, 2 = moderate discomfort, 3 = severe discomfort, 4 = moribund), clinical signs (including depressed, decreased appetite, nasal discharge, ocular discharge, increased respiratory rate/effort, diarrhea, recumbent unable to rise), rectal temperature and respiration rates. Observations were made at 12-h intervals from inoculation on d 0 through d 8, and at 24-hour intervals from d 9 to d 31 by trained observers. Observers were not blinded to the treatment (BRD vs CON) because the BRD-treated heifers were housed separately from all other cattle; however, observers were blinded as to the TI vs CON status of individual heifers.

### Blood collection and processing for complete blood count (CBC), serum, and white blood cells (WBCs)

Whole blood was collected in vacutainer tubes containing no additive for serology and hematological parameters and in vacutainer tubes containing K_2_EDTA (Becton, Dickinson and Company, Franklin Lakes, NJ 07417) for WBC analysis. Vacutainer tubes containing no anticoagulant were centrifuged at 377 × g for 10 min at 4°C (Eppendorf 5804R, Enfield, CT, USA). The separated serum was placed into microcentrifuge tubes and stored at −20°C for serology. Tubes containing K_2_EDTA were centrifuged at 300 × *g* for 10 min at 4°C. For each sample, the buffy coat was transferred into tubes containing 5 mL of ammonium–chloride–potassium lysing buffer (ACK; KD Medical, Columbia, MD, USA, Cat# RGF-3015) and incubated at room temperature for 5 min. The WBC were pelleted by centrifugation at 42 × g for 10 min at 4°C, resuspended in 2.5 mL of ACK, and centrifuged at 300 × *g* for 10 min at 4°C. Following centrifugation in ACK, the supernatant was decanted, and the WBC pellet was resuspended in 1X PBS (pH 7.4), followed by centrifugation at 300 × *g* for 10 min at 4°C. Following centrifugation, the supernatant was decanted, and the WBC pellets were resuspended in 1.0 ml of Trizol reagent (Ambion, Carlsbad, CA). Samples were stored at-80°C until further analysis. Hematological parameters for WBC, red blood cells (RBC), and platelets were determined using the Element HT5 (Heska, Loveland, CO).

### RNA extraction from WBCs for BVDV RTPCR

RNA was extracted from WBC samples to verify the presence of BVDV for time points from d 0 (prior to inoculation) to d 31 post-inoculation from CON-BRD and TI-BRD heifers by methods previously described (Van Campen et al., 2024). The BVDV PCR reaction products were separated on a 2% agarose gel containing Gel Red (Biotium. Fremont, CA, USA, Cat# 41003) and visualized with a Molecular Imager ChemDocXRS+with Image Lab software (Version 6, BioRad, Hercules, CA, USA) RNA extracted from the WBCs from a type1b ncp BVDV-infected calf was included as a positive control for the BVDV RTPCR reaction.

### Serology

Serum neutralizing (SN) antibody titers for each serum sample were determined using a microtiter plate format with either the cytopathic (cp) type 1b BVDV (TGAC) or the cp type 2 BVDV (296c) and BTs as indicator cells (Carbrey et al., 1971). Serial 2-fold dilutions of each serum sample were made in duplicate in 96-well plates, and 2.33 log_10_ TCID_50_ of cp type 2 BVDV virus (296c) or 2.5 log_10_ TCID_50_ of cp type 1b BVDV (TGAC) in 25 µl was added to each well. The plates were incubated for 1 hour at 37°C, 5% CO_2_, before adding 1 × 10^4^ BT in 150 µl to each well. The 96-well plates were incubated at 37°C, 5% CO_2_ for an additional 4 days. The BTs in each well were viewed with a phase contrast microscope (Nikon TS100) and scored for cp effect. The highest dilution of serum that prevented cp changes in the BT cells, expressed as the number of 2-fold dilutions of the serum sample, i.e., 2^n^, was recorded as the SN titer. Controls included serum from a calf vaccinated with a modified live viral vaccine containing both type 1 and type 2 BVDV (Bovishield Gold, Zoetis), and wells containing uninfected BTs. The amount of cp BVDV in the inoculum was determined by serial dilution, and the TCID_50_ was calculated (Reed and Muench, 1938).

### Flow cytometry

Staining for immune cell markers and flow cytometry were performed on WBCs as previously described (Kincade et al., 2025). Whole blood was collected in K2EDTA tubes. Red blood cells (RBCs) were lysed with 1x RBC lysis buffer (eBioscience, San Diego, CA, USA, cat# 00-4300-54). The samples were centrifuged and washed multiple times with 1x PBS. The remaining cell pellet contained white blood cells (WBCs), which were then plated in 96 well plates and stained. The WBCs were stained with 100 µL of Fixable Viability Stain 620 (BD Biosciences, Franklin Lakes, NJ, USA, cat# 564996) and a master mix of labeled antibodies and analyzed at the CSU Flow Cytometry Core on a 4L 16V-14B-10YG-8R Cytek Aurora spectral cytometer. The flow cytometry data were gated using FlowJo (BD Biosciences, Franklin Lakes, NJ, USA), and the frequency of each subpopulation was expressed relative to its parent population (Kincade et al., 2025).

### Statistical analysis

All statistical analyses were conducted in R (R Core Team, 2023) using RStudio software. Outlier tests were performed on all data (± 3 STDEV from the mean), and no outliers were removed from the data set. Final body weight and average daily gain were assessed using a linear model, with TI, BRD, and their interaction as the fixed effect, and initial body weight as a covariate. Hematological data were assessed using a restricted maximum likelihood-based, mixed-effects model, and repeated-measures (repeated blood samples and body weight (BW) analysis. TI (CON or TI), BRD (CON or BRD), day post BVDV inoculation, and their interaction were included in the model as fixed effects. The animal was considered the experimental unit for all response variables measured, which was accounted for as a random variable in the mixed effects model when applicable. A type 3 ANOVA table was constructed using the Kenward–Roger method of computing denominator degrees of freedom. Significance was determined at P ≤ 0.05, and tendencies were determined if P > 0.05 and ≤ 0.10. When a significant three-way interaction was observed, least squares means were compared within that interaction. In the absence of a three-way interaction, two-way interactions or main effects were evaluated accordingly. Due to the high number of total variables evaluated, results were grouped according to the highest-order significant interaction, and only significant results were presented to improve presentation and reduce table complexity.

## RESULTS AND DISCUSSION

Bovine viral diarrhea viruses are enveloped, single-stranded, positive-sense RNA viruses belonging to the Pestivirus genus in the family Flaviviridae, which infect a variety of ruminant species and swine worldwide. Bovine viral diarrhea viruses are classified as type 1 (Pestivirus A) and type 2 (Pestivirus B; Postler et al., 2023). The majority of BVDV biotypes are non-cytopathic (ncp) in cell culture, with cytopathic (cp) biotypes arising from mutations within multiple coding regions in the viral genome (Tautz et al., 2015). Acute BVDV infections result in a wide range of clinical presentations from inapparent to severe, depending on the virus strain (Pellerin et al., 1994; Bielefeldt-Ohmann, 1995). Earlier studies have revealed that TI cattle had impaired performance (Engle et al., 2024), which could be due to epigenetic dysregulation of the immune system in TI cattle (Van Campen et al., 2024; Kincade et al., 2025). As TI calves make up the larger proportion of calves in BVDV-infected herds, in comparison to persistently infected calves (Muñoz-Zanzi et al., 2003), the reduction in growth, health, and increased susceptibility to disease may represent a greater share of the economic cost of fetal BVDV infections.

Along with BVDV infection, BRD is also one of the major issues in the cattle industry, and the link between TI and susceptibility and/or resistance to BRD remains underexplored. In the current study, TI heifers were generated and were inoculated with type 1b BVDV and *M. haemolytica* to investigate the link between fetal TI and potential postnatal resistance to BRD. Across the challenge study, a mild increase in the clinical scores (P < 0.05) was observed on days 3 and 4 in the BRD-challenged (d0 and 1 after *M. haemolytica* inoculation) heifers compared to control heifers (Table 3). Clinical signs noted in BRD heifers included nasal discharge, coughing, and diarrhea. Rectal temperatures and respiration rates were not different (P > 0.10) between groups on any day. Only 1 heifer in the CON-BRD group met the criterion (rectal temperature ≥ 40.6°C) for antibiotic treatment and was further excluded from the study. This animal responded to treatment with a decrease in rectal temperature within 12 h. There was no difference (P > 0.10) in BW or rate of gain between the 4 groups over the 31-day study period.

The number of RBC/µL and the hemoglobin content were similar (P > 0.10) across treatment groups (data not shown). A tendency for a 3-way interaction between TI, BRD, and days was detected for WBC (P = 0.056). Total WBCs were lower (P < 0.05) in CON-BRD heifers compared to CON-CON and TI-CON heifers on days 0, 6, and 8 (Table 1 and Supplemental Figure S1). The leukopenia detected in CON-BRD heifers is due to the decreased numbers of lymphocytes and basophils observed in the CBC analysis of TI and BRD heifers (Table 2 and 3; Supplemental Figures S1). The timing and extent of leukopenia and lymphopenia relative to type 1b ncp BVDV inoculation are similar to the results of experimental acute infections with BVDV on WBCs previously reported (Bolin et al., 1985; Ellis et al., 1988; Gånheim et al., 2005). There was no effect of TI status on neutrophil counts in the current study, which is different from the findings observed by Burciaga-Robles et al (2010), where they observed a decrease in neutrophils in steers 72 hours after exposure to PI steers and an increase in neutrophils following intratracheal *M. haemolytica* challenge. The lymphopenia post-BVDV inoculation in BRD heifers relative to the CON treatment groups is typical of acute BVDV infections (Ellis et al., 1998) and as described in the BRD challenge experiment on which this challenge was based (Burciaga-Robles et al., 2010). The current study also observed a greater (P < 0.05) platelet count in TI heifers compared to the controls (Kocaturk, M. et al., 2010.) In contrast, heifers infected with BRD had a lower (P < 0.05) platelet count compared to the heifers that were not inoculated with BRD. Monocytes and eosinophils were also not impacted (P < 0.10) by TI or BRD as determined from the CBC in the current study.

**Table 1.**
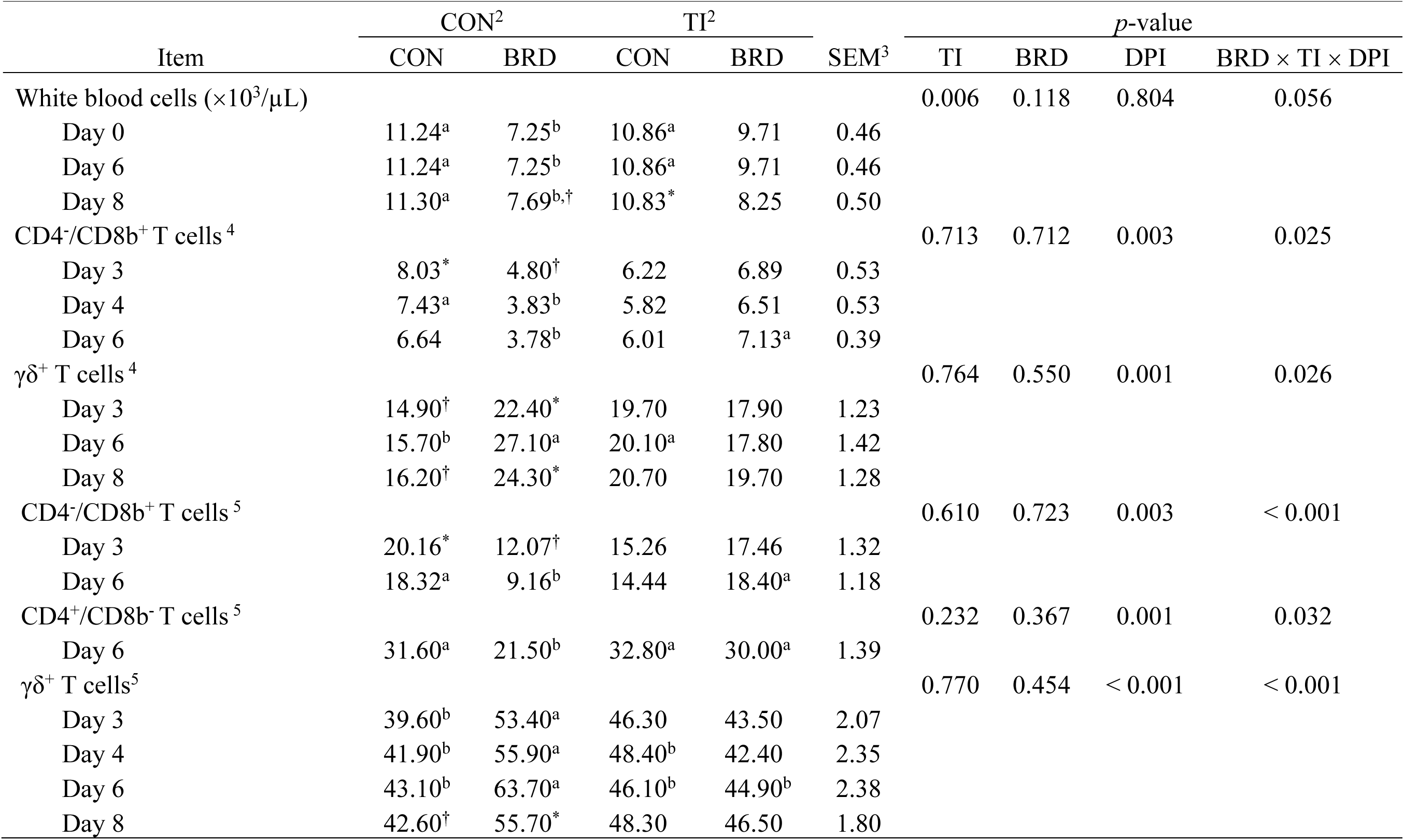
Effects of fetal transient bovine viral diarrhea virus (BVDV) infection (TI), bovine respiratory disease (BRD), day post infection (DPI), and their interactions on hematological measures^1^ in heifers after inoculation with BVDV1b and Mannheimia haemolytica.

|  | CON<sup>2</sup> |  | TI<sup>2</sup> |  |  | *p*-value |  |  |  |
|---|---|---|---|---|---|---|---|---|---|
| Item | CON | BRD | CON | BRD | SEM<sup>3</sup> | TI | BRD | DPI | BRD × TI × DPI |
| White blood cells (×10<sup>3</sup>/µL) |  |  |  |  |  | 0.006 | 0.118 | 0.804 | 0.056 |
| Day 0 | 11.24<sup>a</sup> | 7.25<sup>b</sup> | 10.86<sup>a</sup> | 9.71 | 0.46 |  |  |  |  |
| Day 6 | 11.24<sup>a</sup> | 7.25<sup>b</sup> | 10.86<sup>a</sup> | 9.71 | 0.46 |  |  |  |  |
| Day 8 | 11.30<sup>a</sup> | 7.69<sup>b,†</sup> | 10.83<sup>*</sup> | 8.25 | 0.50 |  |  |  |  |
| CD4<sup>-</sup>/CD8b<sup>+</sup> T cells<sup>4</sup> |  |  |  |  |  | 0.713 | 0.712 | 0.003 | 0.025 |
| Day 3 | 8.03<sup>*</sup> | 4.80<sup>†</sup> | 6.22 | 6.89 | 0.53 |  |  |  |  |
| Day 4 | 7.43<sup>a</sup> | 3.83<sup>b</sup> | 5.82 | 6.51 | 0.53 |  |  |  |  |
| Day 6 | 6.64 | 3.78<sup>b</sup> | 6.01 | 7.13<sup>a</sup> | 0.39 |  |  |  |  |
| γδ<sup>+</sup> T cells<sup>4</sup> |  |  |  |  |  | 0.764 | 0.550 | 0.001 | 0.026 |
| Day 3 | 14.90<sup>†</sup> | 22.40<sup>*</sup> | 19.70 | 17.90 | 1.23 |  |  |  |  |
| Day 6 | 15.70<sup>b</sup> | 27.10<sup>a</sup> | 20.10<sup>a</sup> | 17.80 | 1.42 |  |  |  |  |
| Day 8 | 16.20<sup>†</sup> | 24.30<sup>*</sup> | 20.70 | 19.70 | 1.28 |  |  |  |  |
| CD4<sup>-</sup>/CD8b<sup>+</sup> T cells<sup>5</sup> |  |  |  |  |  | 0.610 | 0.723 | 0.003 | < 0.001 |
| Day 3 | 20.16<sup>*</sup> | 12.07<sup>†</sup> | 15.26 | 17.46 | 1.32 |  |  |  |  |
| Day 6 | 18.32<sup>a</sup> | 9.16<sup>b</sup> | 14.44 | 18.40<sup>a</sup> | 1.18 |  |  |  |  |
| CD4<sup>+</sup>/CD8b<sup>-</sup> T cells<sup>5</sup> |  |  |  |  |  | 0.232 | 0.367 | 0.001 | 0.032 |
| Day 6 | 31.60<sup>a</sup> | 21.50<sup>b</sup> | 32.80<sup>a</sup> | 30.00<sup>a</sup> | 1.39 |  |  |  |  |
| γδ<sup>+</sup> T cells<sup>5</sup> |  |  |  |  |  | 0.770 | 0.454 | < 0.001 | < 0.001 |
| Day 3 | 39.60<sup>b</sup> | 53.40<sup>a</sup> | 46.30 | 43.50 | 2.07 |  |  |  |  |
| Day 4 | 41.90<sup>b</sup> | 55.90<sup>a</sup> | 48.40<sup>b</sup> | 42.40 | 2.35 |  |  |  |  |
| Day 6 | 43.10<sup>b</sup> | 63.70<sup>a</sup> | 46.10<sup>b</sup> | 44.90<sup>b</sup> | 2.38 |  |  |  |  |
| Day 8 | 42.60<sup>†</sup> | 55.70<sup>*</sup> | 48.30 | 46.50 | 1.80 |  |  |  |  |

**Table 2.** Effect of fetal transient bovine viral diarrhea virus (BVDV) infection (TI) status and days post infection (DPI) on hematological variables in heifers after BVDV1b and Manheimia haemolytica challenge. ^1^.

|  | TI<sup>2</sup> |  |  | *p*-value |  |  |
|---|---|---|---|---|---|---|
| Item | Control | TI | SEM<sup>3</sup> | TI | DPI | TI × DPI |
| Platelet (×10<sup>3</sup>/µL) | 375 | 494 | 38 | 0.041 | 0.318 | 0.484 |
| Basophils (×10<sup>3</sup>/µL) | 0.029 | 0.040 | 0.004 | 0.058 | 0.278 | 0.432 |
| BVDV2 SN titer<sup>4</sup> | 70.17 | 0.64 | 1.85 | < 0.001 | 0.132 | 0.902 |

<sup>1</sup>Values are presented as least squares means. All variables were analyzed using a model that included the fixed effects of TI status, BRD status, DPI, and all associated interactions. Due to the non-significant three-way interaction, TI status and DPI interaction were presented.
<sup>2</sup> Status of heifers: Control = Heifers from dams that were inoculated with sham control media. TI = Heifers from dams that were inoculated with noncytopathic type 2 BVDV at gestational day 175.
<sup>3</sup>SEM = standard error of the mean.
<sup>4</sup>BVDV2 SN=noncytopathic type 2 bovine viral diarrhea virus serum neutralizing titers.

**Table 3.** Effect of bovine respiratory disease (BRD) status and days post infection (DPI) on clinical scores and CBC hematological measures^1^ in heifers after inoculation of BVDV1b and Mannheimia haemolytica.

|  | BRD<sup>2</sup> |  |  | p-value |  |  |
|---|---|---|---|---|---|---|
| Item | Control | BRD | SEM<sup>3</sup> | BRD | DPI | BRD × DPI |
| Clinical score |  |  |  | 0.013 | 0.014 | 0.014 |
| Day 3 | 0.00 | 0.28 | 0.28 |  |  |  |
| Day 4 | 0.00 | 0.53 | 0.11 |  |  |  |
| Lymphocytes (×10<sup>3</sup>/µL) | 7.66 | 6.13 | 0.31 | 0.002 | 0.258 | 0.678 |
| Basophils (×10<sup>3</sup>/µL) | 0.041 | 0.028 | 0.004 | 0.029 | 0.278 | 0.615 |
| Platelet (×10<sup>3</sup>/µL) | 537.0 | 332.0 | 38.3 | 0.001 | 0.318 | 0.233 |
| Intermediate Monocytes<sup>4</sup> |  |  |  | 0.100 | 0.060 | 0.001 |
| Day 3 | 1.08 | 1.85 | 0.23 |  |  |  |
| Day 4 | 0.96 | 1.42 | 0.16 |  |  |  |
| Day 6 | 0.91 | 1.69 | 0.17 |  |  |  |
| Day 17 | 1.46 | 0.86 | 0.10 |  |  |  |
| Day 31 | 1.06 | 1.57 | 0.17 |  |  |  |
| CD4<sup>-</sup>/CD8b<sup>-</sup> T cells |  |  |  |  |  |  |
| Day 6 | 5.99 | 4.99 | 0.31 | 0.602 | < 0.001 | 0.002 |

<sup>1</sup>Values are presented as least squares means. All variables were analyzed using a model that included the fixed effects of TI status, BRD status, DPI, and all associated interactions. Due to the non-significant three-way interaction, BRD status and DPI interaction were presented.
<sup>2</sup> BRD = Status of heifers: Control = heifers received 2 ml of media (Dulbecco's minimal essential medium; DMEM + 2% horse serum; HS) instilled into each nostril (total of 4 ml) on d 0, followed by intratracheal instillation of 10 ml of phosphate-buffered saline on d 3. BRD = heifers received 2 ml of 4.0 log<sub>10</sub> TCID<sub>50</sub> of the ncp type 1b BVDV in DMEM + 2% HS instilled into each nostril (total of 4 ml) on d 0, followed by intratracheal instillation of *M. haemolytica* on d 3.
<sup>3</sup>SEM = standard error of the mean.
<sup>4</sup>Intermediate monocytes (CD45<sup>+</sup>/CD11b<sup>+</sup>/CD172a<sup>+</sup>/CD14<sup>+</sup>/CD16<sup>+</sup>) are presented as a percentage of total CD45<sup>+</sup> cells.
<sup>5</sup>CD4<sup>-</sup>/CD8b<sup>-</sup> T cells (CD45<sup>+</sup>/CD3<sup>+</sup>/γδ<sup>-</sup>/CD335<sup>-</sup>) are presented as a percentage of total CD3<sup>+</sup> cells.

Viral RNA was detected in WBC RNA by RTPCR from CON-BRD heifers beginning on d3 post-inoculation with type 1b BVDV, and amplicons were detected in all 6 CON-BRD heifers on d 6 and on d 8 post-inoculation. Amplicons were not observed after the appearance of serum neutralizing antibodies (Figure 1). BVDV RNA was not detected in WBCs of TI-BRD heifers, from d0 through d10 post-inoculation with type 1b BVDV.

**Figure 1.**
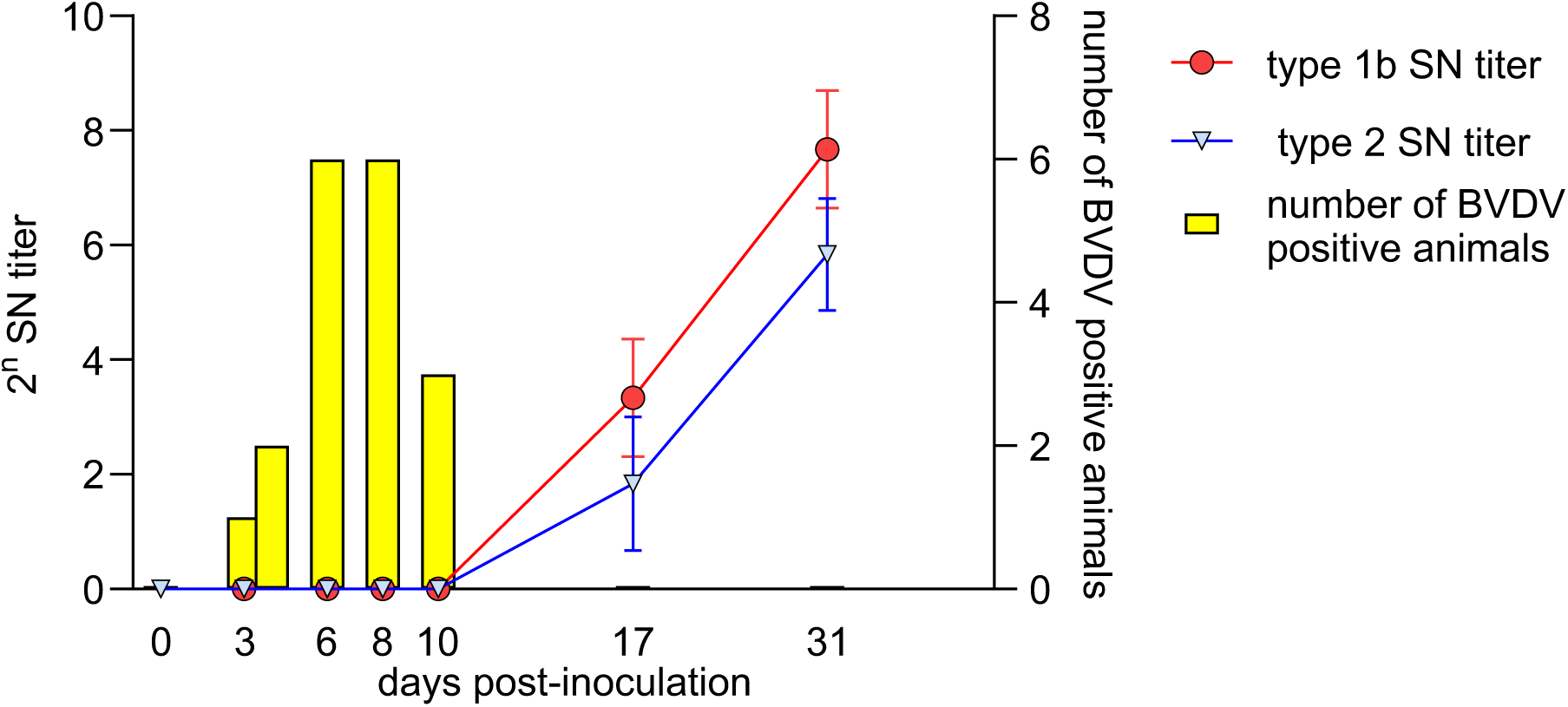
The number of CON-BRD heifers (uninfected heifers received 2 ml of 4.0 log10 TCID50 of the ncp type 1b BVDV in DMEM + 2% HS instilled into each nostril (total of 4 ml) on d 0, followed by intratracheal instillation of M. haemolytica on d 3) in which amplicons were detected by BVDV RTPCR in WBCs on days post-inoculation with type 1b BVDV, compared to the appearance of serum neutralizing antibody titers (2^n^ +/- SEM) to type 1b and to type 2 BVDV.

A 3-way interaction between TI, BRD, and Day was detected (P < 0.05) for the following flow cytometry populations: CD4^-^/CD8b^+^ T cells and γδ^+^ T cells as a percentage of both total CD45^+^ cells and total CD3^+^ cells, and CD4^+^ CD8b^-^T cells as a frequency of total CD3^+^ cells. The CD4^+^/CD8b^-^ (as a percentage of total CD3^+^ cells) were lower (P < 0.05) in the CON-BRD heifers on day 6 compared to CON-CON heifers (Table 1). The CD4^-^/CD8b^+^ cells (as a percentage of either total CD45^+^ or total CD3^+^ cells) were lower (P < 0.05) in CON-BRD heifers compared to CON-CON heifers on day 3, 4, and 6 post challenge (Table 1). The frequency of CD4^-^/CD8b^-^ T cells (as a percentage of total CD3^+^ cells) increased and CD4-/CD8b-T cells (as a percentage of total CD3+ cells) decreased on day 6 compared to controls. Intermediate monocytes (as a percentage of total CD45^+^ cells) were increased on days 3, 4, 6, and 31 compared to control heifers. Intermediate monocytes (as a percentage of total CD45+ cells) decreased in BRD infected heifers on day 17.

Gamma-delta (γδ^+^) T cells increased (as a percentage of total CD45^+^ and total CD3^+^ cells) in CON-BRD heifers compared to CON-CON (on day 3, 4 ( γδ^+^ /CD3^+^ only), 6, and 8), TI-CON (day 4 and 6), and TI-BRD (day 6) heifers (Table 1). Circulating γδ^+^ T cells in cattle are early responders to foreign antigens with cytotoxic (Lundberg and Splitter, 2000; Endsley et al., 2004), antigen-presentation (Collins et al., 1998), and regulatory functions including the release of cytokines, IL-17 (McGill et al., 2016), IFNγ, TGFβ (Rhodes et al., 2001), and IL-10 (Hoek et al., 2009). Thus, the increase in γδ^+^ T cells observed in this experimental BVDV inoculation is an expected response to a primary viral infection in a naïve animal. B-cells, CD4^-^/CD8b^-^ T cells, CD4^+^/CD8b^-^ T cells, CD4^+^/CD8b^-^/CD25^+^/CD127^-^ cells, natural killer T cells, classical monocytes, granulocytes, and non-classical monocytes, (as a percentage of total CD45^+^), in addition to CD4^+^/CD8b^+^ T cells and natural killer T cells (as a percentage of total CD3^+^ cells), were not impacted by any of the variables tested (P > 0.10; TI, BRD, Day, or their interactions).

TI heifers had greater (P < 0.01) SN titers to type 2 BVDV (2^n^ = 70) compared to CON heifers (2^n^ = 0.64), including before BVDV inoculation. TI heifers also had a slightly greater (P < 0.01) cross-reactive SN titer to type 1b BVDV (4.7) compared to CON heifers (0.91; CON-CON and CON-BRD). TI-CON and TI-BRD heifers maintained stable SN titers to type 2 BVDV over the 31-day period (data not shown). TI heifers (TI-CON and TI-BRD) had a greater (P < 0.05) BVDV1b SN titer compared to CON heifers on days 0, 3, 6, and 10. CON heifers (CON-CON and CON-BRD) had similar (titer = 0) BVDV1b titer from day 0 to day 10. Serum neutralizing antibodies to type 1b BVDV were first detected in CON-BRD serum samples on d 17 post-BVDV inoculation, which was greater (P < 0.05) than observed in CON-CON. Type 1b BVDV antibodies in CON-BRD increased by > 2-fold by d 31, which was greater (P < 0.05) than CON-CON and TI-CON. Similar to the CON group, TI heifers (TI-CON and TI-BRD) had similar BVDV1b titers from d 0 to 10, and starting on d 17 until 31, BVDV1b titer in TI-BRD heifers increased slightly and was greater (P < 0.05) than the BVDV1b titer in TI-CON heifers. BRD challenge increased the BVDV1b titers in CON-BRD and TI-BRD heifers starting on day 17, but CON-BRD heifers had a greater increase compared to TI-BRD heifers. The appearance of SN antibodies was delayed compared to the time course for seroconversion reported by Burciaga-Robles et al. (2010), in which type 1b BVDV SN titers were first detected on d 7 following exposure to PI heifers for 72 h. CON-CON heifers remained seronegative to both type 1b (data not shown) and type 2 (Figure 2) BVDVs throughout the experimental period in the current study, as they were purposefully not infected with BVDV.

**Figure 2.**
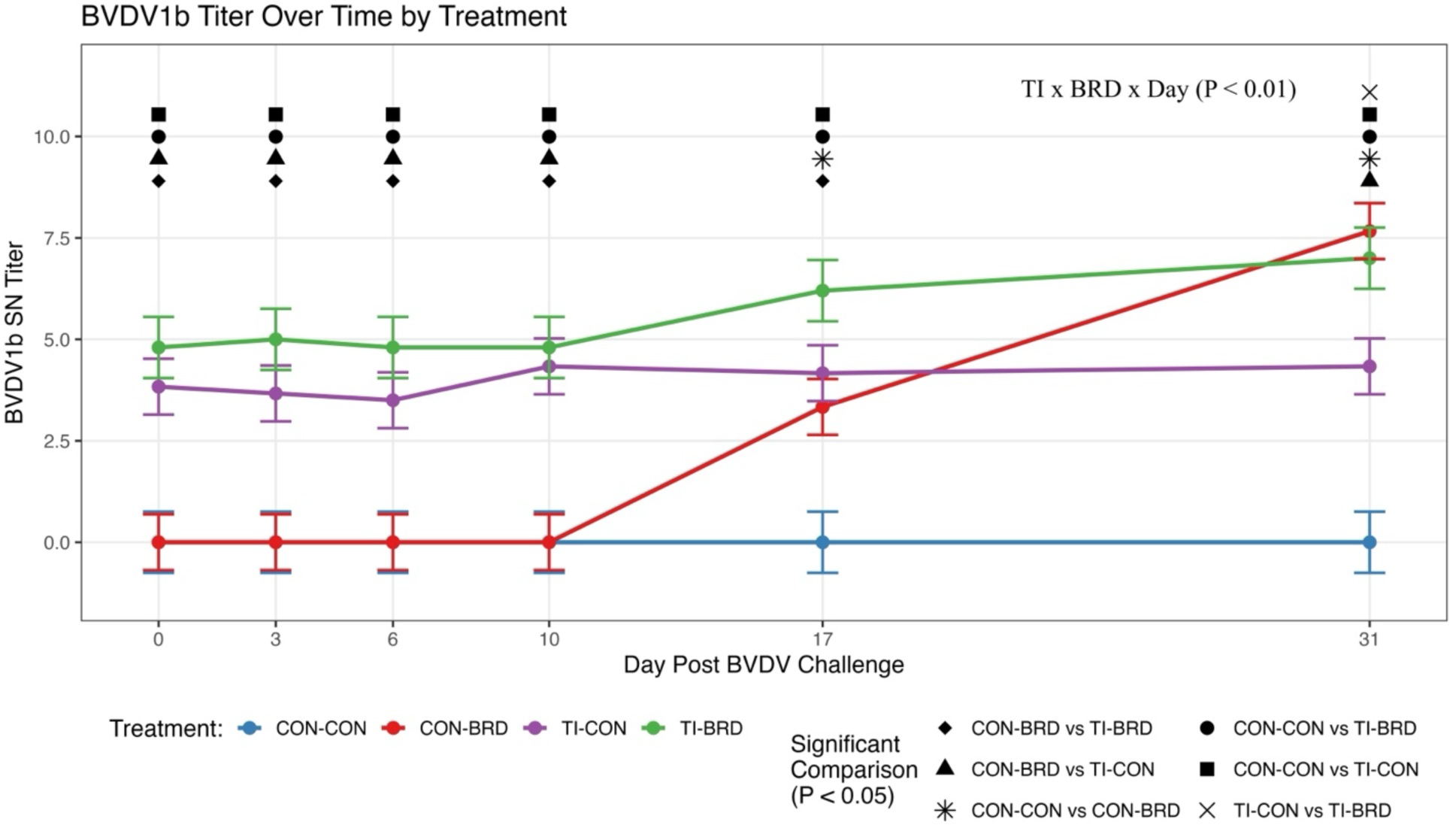
Type 1b bovine viral diarrhea virus (BVDV) SN titer in heifers over 31 days of BVDV and *Mannheimia haemolytica* challenge. Treatments are: 1) CON-CON: uninfected heifers received 2 ml of media (Dulbecco’s minimal essential medium; DMEM + 2% horse serum; HS) instilled into each nostril (total of 4 ml) on d 0, followed by intratracheal instillation of 10 ml of phosphate-buffered saline (PBS) on d 3 (n = 5), 2) TI – CON: TI heifers received 2 ml of media (DMEM + 2% HS) instilled into each nostril (total of 4 ml) on d 0, followed by intratracheal instillation of 10 ml of PBS on d 3 (n=6), 3) CON – BRD: uninfected heifers received 2 ml of 4.0 log10 TCID50 of the ncp type 1b BVDV in DMEM + 2% HS instilled into each nostril (total of 4 ml) on d 0, followed by intratracheal instillation of M. haemolytica on d 3 (n = 6), and 4) TI – BRD: TI heifers received 2 ml of 4.0 log10 TCID50 of the ncp type 1b BVDV in DMEM + 2% HS instilled into each nostril (total of 4 ml) on d 0, followed by intratracheal instillation of M. haemolytica on d 3 (n = 5). Different symbols indicate the significant comparison between different treatment pairs.

There were important differences between the BRD challenge used in this experiment and the Burciaga-Robles et al. (2010) study, which may explain the mild clinical response of the heifers challenged in this experiment. Unlike the previous study, the heifers were acclimated to their pens and were not subjected to the stresses of transportation, adjustment to a new location, feed, or social stress prior to inoculation with BVDV and *M. haemolytica*. The type 1b ncp BVDV inoculum was administered in a single dose by intranasal instillation as opposed to exposure to PI cattle for 72 h, likely reducing the amount of BVDV received by each heifer. Additionally, the intranasal inoculation may have limited the initial BVDV infection to the nasopharynx; whereas the size of particles containing the type 1b BVDV aerosolized in respiratory secretions from PI animals potentially delivered the virus to a greater depth in the respiratory tract (Hatch, 1961). Nonetheless, in this study, the challenged cattle were infected via the intranasal inoculation of BVDV as evidenced by the detection of viremia, lymphopenia, and seroconversion to type 1b BVDV. *M. haemolytica* is a commensal of the nasopharynx of cattle (Klima et al., 2011). Nasopharyngeal samples obtained at 4 and 7 months of age detected *M. haemolytica* DNA in 15 of the 22 heifers. It is likely that these heifers had some adaptation or resistance to *M. haemolytica* prior to inoculation as a part of their symbiotic relationship.

## Conclusion

Type 2 BVDV antibodies acquired during fetal infection during late gestation provided cross-protective immunity that reduced the severity of clinical signs, viremia, leukopenia, lymphopenia, and thrombocytopenia in 8-month-old TI heifers following challenge with a ncp type 1b BVDV and *M. haemolytica*. This study indicates that the prior specific immunity to BVDV resulting from fetal transient BVDV infection conferred protection against lymphopenia, viremia, and clinical disease in TI heifers following postnatal challenge with a combination of a ncp type 1b BVDV and M. haemolytica. Although immune system dysfunction in TI animals was observed in a previous study, evidence for immune dysfunction was not observed in this specific experimental challenge model. However, unfavorable outcomes of other microbial infections due to altered immune responses in TI cattle may be possible.

## ACKNOWLEDGEMENTS

The authors would like to extend thanks to the following personnel: Aydin Guzeloglu (ARBL, Colorado State University), Whitney Simmons, Christina Nash, and Micah Schmidt (ARDEC, Colorado State University), Michael Russell and Josh Daniels (VDL Bacteriology Section, Colorado State University) for the *Mannheimia haemolytica* isolate, Erlene Chek and Harrison Hallmark (Department of Animal Science, Colorado State University).

## List of abbreviations

ADG: average daily gain;
BRD: bovine respiratory disease;
BRDC: bovine respiratory disease complex;
BT: bovine turbinate cells;
BVDV: bovine viral diarrhea virus;
BW: body weight;
CBC: complete blood count;
cp: cytopathic;
d: day;
DMEM: Dulbecco’s minimal essential medium;
ncp: noncytopathic;
HS: horse serum;
PBS: phosphate buffered saline;
p.i.: post-inoculation;
RTPCR: reverse transcriptase polymerase chain reaction;
SEM: standard error of the mean;
SN: serum neutralization;
TCID_50_: tissue culture infectious dose 50;
TI: transient fetal infection,

